# Monod law is insufficient to explain biomass growth in nitrogen-limited yeast fermentation

**DOI:** 10.1101/2021.01.05.425518

**Authors:** David Henriques, Eva Balsa-Canto

**Affiliations:** Bioprocess engineering group, IIM-CSIC, Vigo, Spain

**Author notes:** Address correspondence to David Henriques.

**Keywords:** problematic fermentations, nitrogen-limited, kinetic model, *S. cerevisisae*, biomass, wine, sluggish

## Abstract

The yeast *Saccharomyces cerevisiae* is an essential microorganism in food biotechnology; particularly, in wine and beer making. During wine fermentation, yeasts transform sugars present in the grape juice into ethanol and carbon dioxide. The process occurs in batch conditions and is, for the most part, an anaerobic process. Previous studies linked limited-nitrogen conditions with problematic fermentations, with negative consequences for the performance of the process and the quality of the final product. It is, therefore, of the highest interest to anticipate such problems through mathematical models. Here we propose a model to explain problematic fermentations under nitrogen-limited anaerobic conditions. We separated the biomass formation into two phases: growth and carbohydrate accumulation. Growth was modelled using the well-known Monod law while carbohydrate accumulation was modelled by an empirical function, analogous to a proportional controller activated by the limitation of available nitrogen. We also proposed to formulate the fermentation rate as a function of the total protein content when relevant data are available. The final model was used to successfully explain a series of experiments taken from the literature, performed under normal and nitrogen-limited conditions. Our results revealed that Monod law is insufficient to explain biomass formation kinetics in nitrogen-limited fermentations of *S. cerevisiae*. The goodness-of-fit of the herewith proposed model is superior to that of previously published models, offering the means to predict, and thus control, problematic fermentations.

**IMPORTANCE:** Problematic fermentations still occur in the winemaking industrial practise. Problems include sluggish rates of fermentation, which have been linked to insufficient levels of assimilable nitrogen. Data and relevant models can help anticipate poor fermentation performance. In this work, we proposed a model to predict biomass growth and fermentation rate under nitrogen-limited conditions and tested its performance with previously published experimental data. Our results show that the well-known Monod law does not suffice to explain biomass formation. A second term accounting for carbohydrate accumulation is required to predict successfully, and thus control, problematic fermentations.

## INTRODUCTION

The yeast species *Saccharomyces cerevisiae* is the best-studied eukaryote. *S. cerevisiae* presents unique characteristics such as its fermentation capacity and its ability to withstand adverse conditions of osmolarity and low pH. These desirable properties have made of *S. cerevisiae* one of the workhorses of bio-industries including food, beverage-especially wine- and biofuel production industries (1).

In winemaking, the fermentation turns grape must into an alcoholic beverage. Wine production occurs in a closed system (i.e. in batch conditions) and is, for the most part, an anaerobic process. During fermentation, yeasts transform sugars present in the must into ethanol and carbon dioxide. A complete alcoholic fermentation is achieved when the residual fermentable sugar is less than 2 g/L. Despite improvements in fermentation control, problematic fermentations such as, stuck - with a higher than desired sugar residual- and sluggish-unusually long-fermentations still occur in real practice.

Various studies have shown that insufficient levels of assimilable nitrogen contribute to stuck or sluggish fermentations (2). A minimum of 120–140 mg/L of assimilable nitrogen is required to achieve a standard fermentation rate, while nitrogen content in grape juice may be as low as 60 mg/L (3). Therefore, it has become common practice to supplement nitrogen-deficient musts with diammonium phosphate (4). Nevertheless, both excessive or insufficient nitrogen can lead to the production of undesired metabolites affecting the organoleptic properties of wine (5, 4).

The need to predict and control wine fermentation has motivated a quest for mechanistic models of alcoholic fermentation (see the reviews by Marín (6) and Miller and Block (7). Several approaches linking nitrogen consumption to growth and fermentation rate have been proposed in the literature. Alternatives differ on the way they explain biomass formation and the relation of cell mass (or sometimes cell numbers) to fermentation rate.

Dynamic kinetic models incorporate the Monod equation to describe biomass growth (8, 9, 10). The estimation of the Monod parameters requires nitrogen and sugars uptake data throughout the fermentation. Otherwise, lack of identifiability of the parameters will limit the predictive capability of the model, and thus logistic functions may be more suitable (11, 12). Both formulations, Monod or logistic, achieve a maximum biomass value which can not be exceeded.

However, Schulze et al. (13) showed that, for nitrogen-limited anaerobic fermentations of *S. cerevisiae*, there is a substantial increment in biomass content after the depletion of ammonia. Besides, measurements of protein and messenger RNA (mRNA) contents revealed that their concentrations stay stable while the concentration of carbohydrates increases. Similar effects have been detected in baker’s yeast exposed to nitrogen starvation but not in the carbon starved cells; see (14) for a fed-batch aerobic example, or (15) for a chemostat anaerobic example. These data would imply that, at least for ammonium deficient scenarios, the use of the Monod model would be insufficient.

Similarly, Varela et al. (16) showed that protein/carbohydrate fractions in the biomass of *S. cerevisiae* vary substantially throughout the fermentation of wine fermentations. To account for this dynamics, Pizarro et al. (17) and Vargas et al. (18) included, in their flux balance analysis models, an empirical function that controls the concentration of carbohydrates in the newly formed biomass as a function of extracellular sugar.

In what regards to the modelling of fermentation rate, previous studies proposed two alternatives. The first implies that the fermentation rate is proportional to biomass or cell numbers (8, 19, 9, 10, 12). The second includes the role of hexose transporters as a separate entity that is dependent on the transport of ammonia (20, 11, 21, 22, 23).

The advantage of the second over the first is that it enables the simulation of the effect of nitrogen additions. Still, further experimental analysis of the underlying hypothesis is required. Also, a good adjustment to the data comes at the cost of a large number of estimated parameters.

Pizarro et al. (17) and Vargas et al. (18) proposed a third alternative. These authors assumed that cells use different hexose transporters at different fermentation stages based on previously published experimental data (24). However, this approach also requires the adjustment of the maximum transport rate for each hexose carrier. As the maximum transport rate is deemed to be a function of temperature and the initial nitrogen concentration, the modelling also involves the estimation of a substantial number of parameters.

In this work, we propose a dynamic kinetic model capable of successfully explaining nitrogen-limited fermentations. The model describes growth and fermentation rate while distinguishing protein from carbohydrate fractions of biomass. The Monod equation describes the growth, and the carbohydrate accumulation observed after the depletion of ammonia is described by an empirical function, analogous to a proportional controller.

We tested the model properties by fitting the data provided by Schulze et al. (13) and Coleman et al. (9) who explored fermentations under various nitrogen regimes. Our results show why standard Monod models often produce poor results in modelling nitrogen-limited fermentations of *S. cerevisiae*.

Fermentation rate can be a function of the biomass or, as proposed here, a function of the protein fraction since the glycolytic enzymes are an important part of the protein pool. Remarkably, even if protein content is not measured, the model is identifiable provided the uptake of hexoses and nitrogen, as well as the production of ethanol, are measured over time.

We compared the performance of the proposed model with the Monod model and the model proposed by Cramer et al. (8) who used it to explain nitrogen-limited fermentations. Our results showed that the proposed model captures the data more accurately than previously published alternatives (up to 6.8 times improvement in normalised root-mean-square error in the fitting of biomass dynamics, and up to 3.8 times improvement in the fitting of the overall process dynamics). In consequence, our model can be used to predict, and thus, control, fermentation problems.

## RESULTS

### Modeling biomass formation during nitrogen-limited fermentation

The microbial growth rate was described as the sum of two different terms. The first, *µ*_*N*_, corresponds to the healthy cell division, mainly when assimilable nitrogen sources (YAN) are abundant. The second, *µ*_*C*_, corresponds to a secondary increase in biomass after the depletion of nitrogen sources-which we assumed corresponds, mostly t°Carbohydrate accumulation. The biomass growth equation reads:

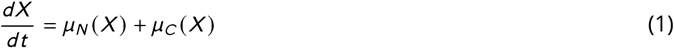

where X is the biomass (g/L).

To model growth associated with cell division we used a Monod type kinetics (25) where nitrogen sources were considered the limiting nutrient:

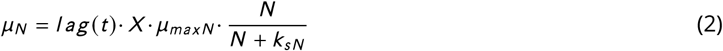

where 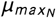 is the maximum specific growth rate, *N* is extracellular YAN (g/L), *k*_*sN*_ is the Monod equation parameter and lag is a function of time (t) representing the lag phase. The lag phase is typically a period of adjustment in which a given microbial population adapts to a new medium before it starts growing exponentially. To model the lag we used the model proposed by (26):

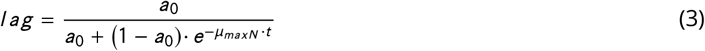

where *a*_0_ is an estimated parameter and *µ*_*maxN*_ is the maximum growth rate associated with *µ*_*N*_ .

The secondary increase in biomass concentration (*µ*_*C*_) is activated by the decrease the concentration of YAN:

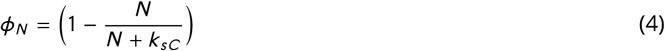

where *k*_*sN*_ is a parameter controlling the half-maximal inactivation of *µ*_*N*_ which is modeled with an empirical expression analogous to a proportional controller:

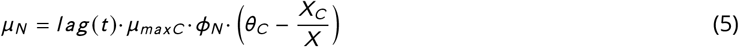

where *θ*_*C*_ is the set-point (target value) for the ratio of carbohydrates in the biomass content (*X*_*C*_ /*X*) and *µ*_*maxC*_ controls the velocity of the convergence towards that reference point.

The generation of carbohydrates is affected both by *µ*_*N*_ and *µ*_*C*_ and its dynamics is described as:

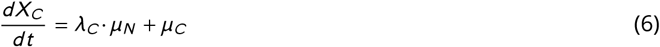

where *λ*_*C*_ is the carbohydrate content in the biomass during exponential growth.

On the contrary the formation of protein is only affected by *µ*_*N*_ and its dynamics is described by:

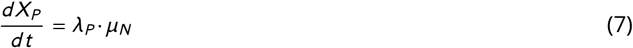

where *λ*_*P*_ is the protein content in the biomass during exponential growth.

YAN is the limiting nutrient in the primary growth phase and its consumption is proportional to *µ*_*N*_ :

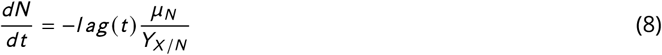

where *Y*_*X* /*N*_ is the YAN to biomass yield.

For the sake of comparison we used two additional models: the first, regarded as *M*_*M*_, corresponds to that for which the biomass follows the Monod description:

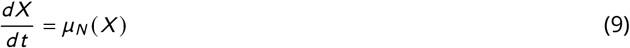

and the second, regarded as *M*_*C*_, corresponds to the model proposed by (8) and subsequently used by (9) to explain nitrogen-limited fermentations. In this model, the biomass follows a Monod equation 9 depending on the active biomass and the dynamics of cell mass includes a death rate proportional to the extracellular ethanol concentration.

### Modeling fermentation rate and production of extracellular metabolites

Regarding the fermentation rate, we proposed two models with different underlying hypotheses:

- M_*P*_: assuming that fermentation rate is proportional to the total protein content (*X*_*P*_),
- M_*X*_: assuming that fermentation rate is proportional to the total biomass content. The uptake of glucose reads as follows:

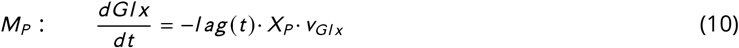

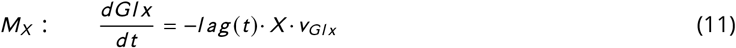

where *v*_*Gl x*_ is the expression governing glucose transport.

Similarly to (27), we used Micheaelis-Menten type kinetics kinetics coupled to ethanol inhibition to model glucose transport:

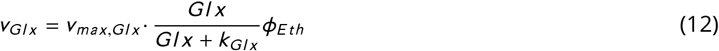

where *v*_*max,Gl x*_ is the maximum rate of glucose transport, *k*_*Gl x*_ is the Michaelis-Menten constant and *φ*_*E th*_ is competitive inhibition induced by ethanol (28) :

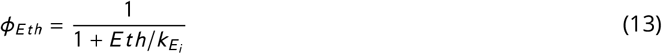

where *E th* is the extracellular ethanol concentration and 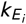 defines the strength of the inhibitory effect.

The excretion of extracellular metabolites was assumed to be proportional to the consumption of glucose. In the case of ethanol, this is described by:

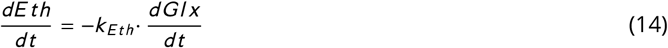

where *k*_*E th*_ controls the quantity of consumed glucose which is converted into ethanol.

### Calibration of candidate models

The Table 1 summarizes the main characteristics of the four candidate models.

**TABLE 1.** Main characteristics of candidate models.

| Model identifier | Model characteristics |
| --- | --- |
| $M_P$ | Biomass is split into two terms, due to nitrogen and carbohydrates<br>Lag-phase<br>Fermentation rate is proportional to the total protein content<br>12 unknown parameters |
| $M_X$ | Biomass is split into two terms, due to nitrogen and carbohydrates<br>Lag-phase<br>Fermentation rate is proportional to the biomass<br>12 unknown parameters |
| $M_M$ | Biomass follows Monod, nitrogen the limiting substrate<br>Lag-phase<br>Fermentation rate is proportional to the total protein content<br>8 unknown parameters |
| $M_C$ | Biomass is split into total and active<br>Lag-phase (not included in the original formulation)<br>Fermentation rate is proportional to the active biomass<br>8 unknown parameters |

The structural identifiability analysis of the models revealed that it is possible to uniquely estimate parameter values provided the dynamics of biomass, glucose uptake, nitrogen sources uptake and ethanol are measured throughout the fermentation. Remarkably, it is not necessary to measure protein for the identification of the model *M*_*P*_.

Candidate models were calibrated by data fitting using data-sets available in the literature. In particular we used data from Schulze et al. (13) and Coleman et al. (9). The protein and carbohydrate contents were estimated from the data provided by Schulze et al. (13) and kept fixed during parameter estimation.

We performed 10 runs of the parameter estimation problem for each case, so to guarantee convergence to the best possible solution. Since we used a metaheuristic for the optimisation of parameters, we obtained a distribution of values. We selected the best fit for ulterior analysis. Table 2 presents the value of the corrected Akaike information criterion for each experiment and model. Results show that, on average, the models proposed in this work, *M*_*P*_ and *M*_*X*_, are significantly better than existing models based on the Monod equation. Only in experiments 4 and 5 models *M*_*P*_, *M*_*X*_ and *M*_*M*_ can not really be distinguished, as their re-scaled Akaike Δ values are within 4 units difference.

**TABLE 2.**
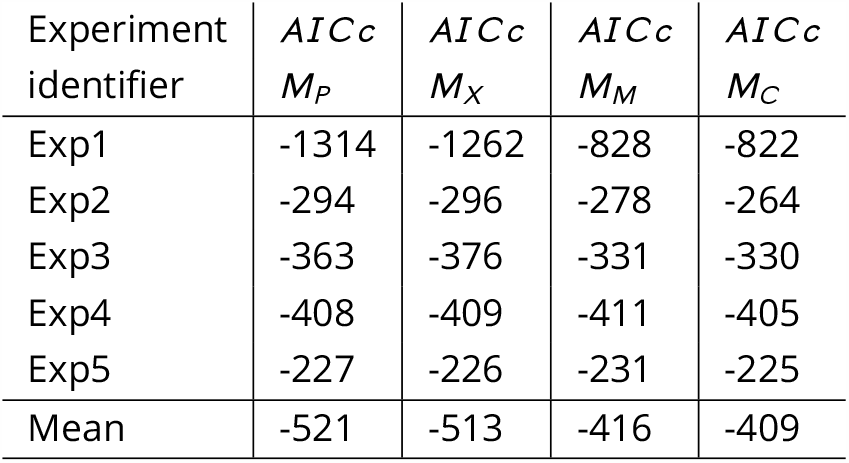
Corrected Akaike information criterion as computed for the different models and experiments.

| Experiment identifier | $AIC_c$<br>$M_P$ | $AIC_c$<br>$M_X$ | $AIC_c$<br>$M_M$ | $AIC_c$<br>$M_C$ |
| --- | --- | --- | --- | --- |
| Exp1 | -1314 | -1262 | -828 | -822 |
| Exp2 | -294 | -296 | -278 | -264 |
| Exp3 | -363 | -376 | -331 | -330 |
| Exp4 | -408 | -409 | -411 | -405 |
| Exp5 | -227 | -226 | -231 | -225 |
| Mean | -521 | -513 | -416 | -409 |

Since the value obtained for the corrected Akaike information does not provide a measure of goodness-of-fit, Figure 1 presents the quality of fit of the different models for the available experiments in terms of the normalised root mean squared error (NMRSE). Figure 1 shows that all models could explain the data within a 10% normalised root mean squared error. However, the performance of the different models differed substantially. Models proposed in this work, *M*_*P*_ and *M*_*X*_, which explain biomass growth using two terms due to nitrogen and carbohydrates, were the best in all experiments; *M*_*P*_ outperforms *M*_*X*_ in 3 out of 5 experiments. NMSRE values for model *M*_*P*_ were within the range 2.42% and 5.72% (obtained for the first and second experiments, respectively); for the model *M*_*X*_, the range was 2.69% to 5.86% (obtained for the first and second experiments, respectively); for model *M*_*M*_ the range was 3.21% and 8.53% (obtained for the fourth and the first experiments respectively); for model *M*_*C*_ the range was 3.49% and 9.32% (obtained for the fourth and the first experiments respectively).

**FIG 1.**
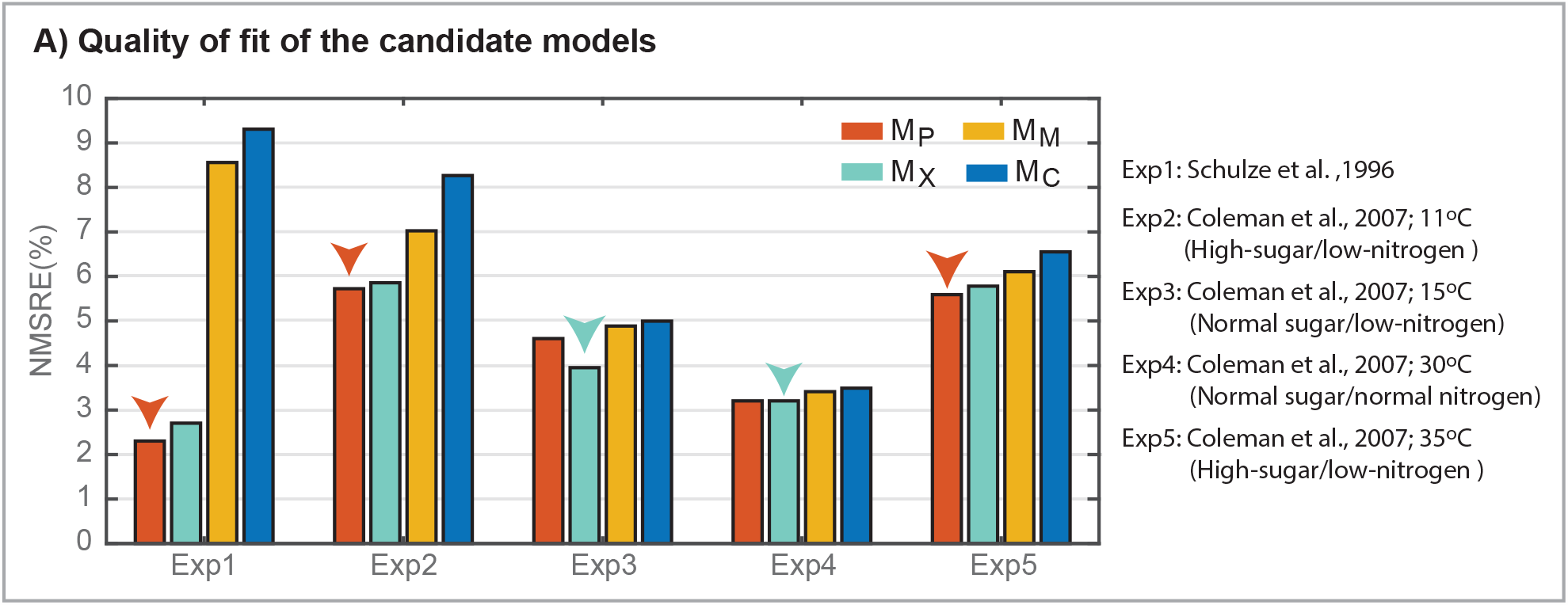
Comparison of candidate models. A) Normalised root-mean-square error for candidate models and all experiments considered. The models proposed in this work, which consider biomass growth split in two phases, due to nitrogen and carbohydrates, are the best in all experiments.

### Biomass formation

Our results suggest that Monod law is insufficient to explain biomass formation kinetics in nitrogen-limited fermentations of *S. cerevisisae*.

Figure 2.A) presents the normalised mean squared error obtained for the biomass for all data sets and models *M*_*P*_ (proposed here), *M*_*M*_ (Monod) and *M*_*C*_ ((9)). Results illustrate the proposed model *M*_*P*_ represents better the data in all cases, even in normal nitrogen conditions. We found the most important difference in the quality of fitting for the (13) data. Remarkably the normalised root-mean-square error obtained for the proposed model (*N M SRE* (*M*_*P*_): 3.04%) is 7.1 times lower than the one obtained with the Monod model (*N M SRE* (*M*_*M*_): 21.49%) and 6.6 times lower than that obtained with the model by Coleman et al. (*N M SRE* (*M*_*C*_): 20.11%). For the experiments in (9), differences are lower; still, the model proposed in this work was always superior in the quality of fit to the biomass data. The improvement was more notorious in those cases using low nitrogen media. Figure 2.B) presents the fit for three out of five experiments. In all three cases, the proposed model could recover the increase of biomass produced after YAN depletion. However, in experiment 5, performed at 35^*o*^ *C* and with high-sugar and low-ethanol, a slight decrease of the biomass is observed in the final data; the model tries to fit the tendency up but also that final data; thus it performs only slightly better than the Monod model.

**FIG 2.**
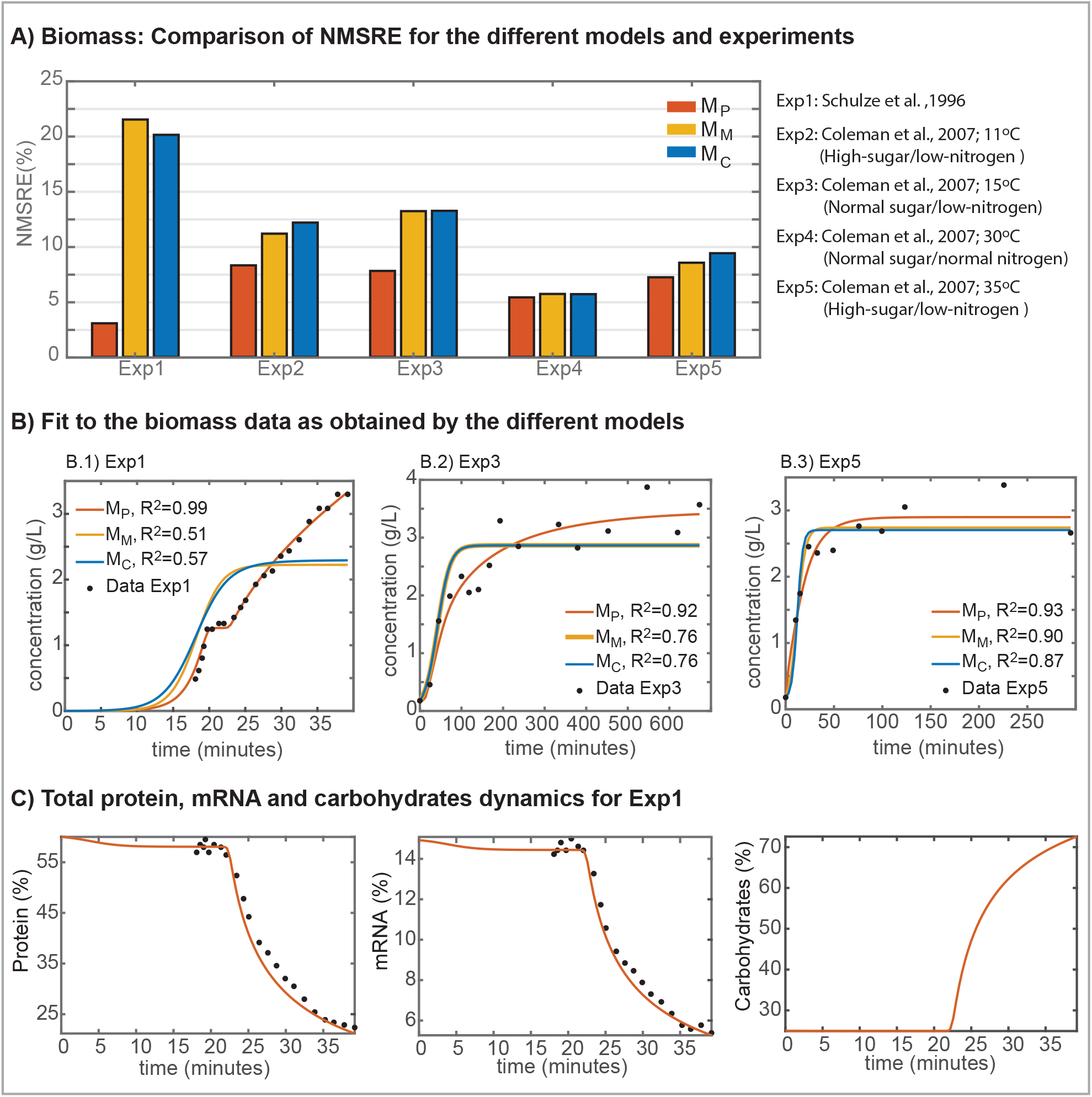
Comparison of candidate models in the fitting of biomass data. A) Normalised root-mean-square error for candidate models and all experiments considered. B) Illustrative examples of the various fit to the data. c) Prediction of the total protein, mRNA and carbohydrates percentage for experiment 1 (in which data are available). Model *M*_*P*_, proposed here, performs better in explaining the biomass as measured by the NMSRE. The model can recover the increase of biomass observed in several experiments after YAN depletion. This increase is significant in experiment 1; while in experiment 3, the increase is followed by a final decrease which is not recovered by the model. Note, however, that the quality of fit is still better even for that particular experiment.

Figure 2.C) shows how after nitrogen depletion, at around 20 min (see Figure 3.A), carbohydrates accumulate and would explain the role of *µ*_2_ in models *M*_*P*_ and *M*_*X*_ in recovering the secondary growth observed in the biomass dynamics.

**FIG 3.**
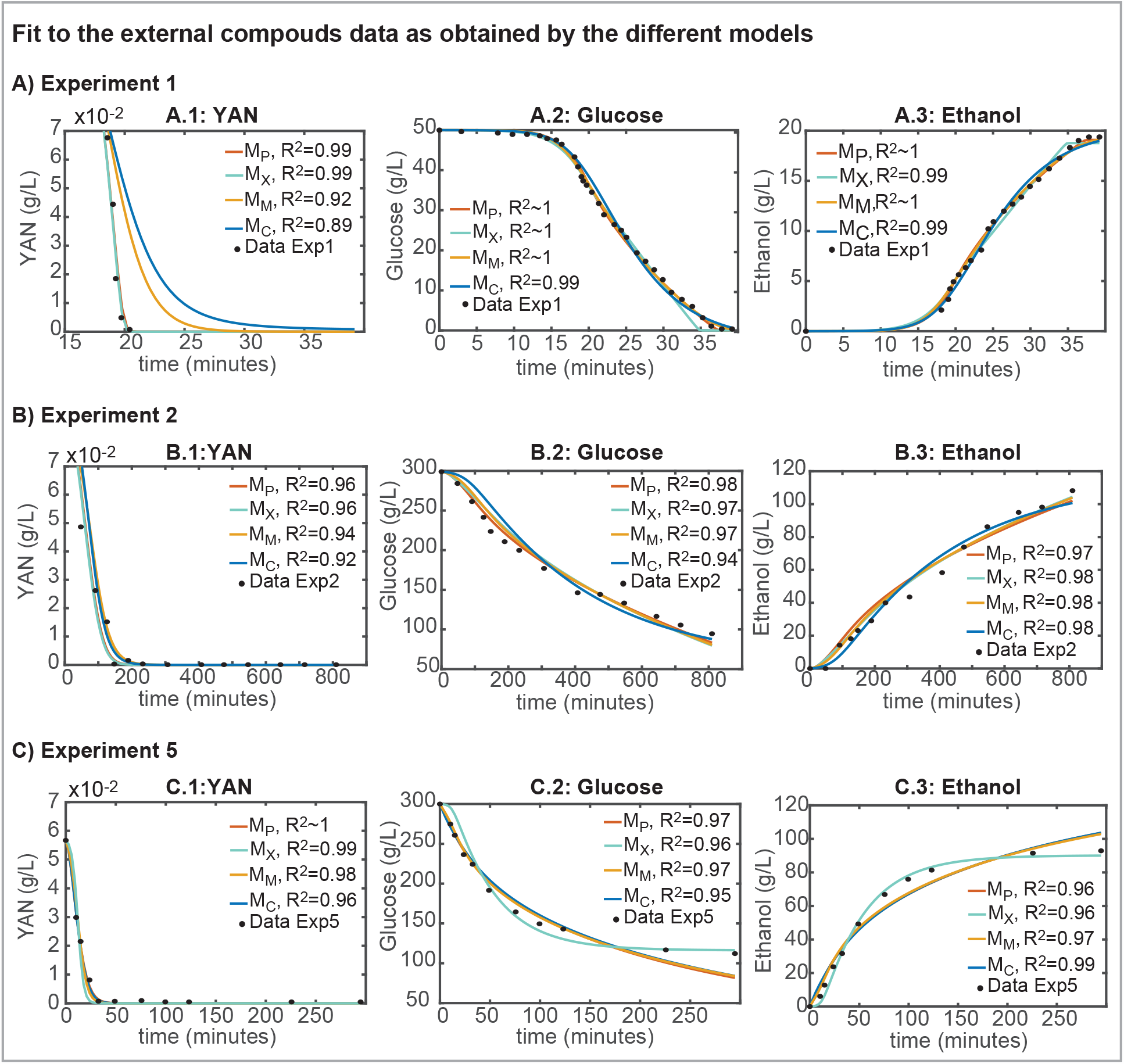
Comparison of candidate models in the fitting of external compounds. A) Experiment 1; B) Experiment 2 and C) Experiment 5. *R*^2^ values are reported for each observable. Note that all models were quite successful in fitting the experimental data. Models *M*_*M*_ and *M*_*C*_ experienced some problems in fitting YAN data; while all models, but *M*_*C*_, found some problems in fitting the data corresponding to a stuck fermentation.

### Fermentation rate

Both models, *M*_*P*_ and *M*_*X*_, were able to recover the overall dynamics of YAN and glucose consumption, and ethanol formation. Remarkably the performance of the model *M*_*X*_, which uses the biomass instead of the protein to explain the fermentation rate is slightly worse than the performance of the model *M*_*P*_ but for the experiment 3 in which normal sugar and low nitrogen were used and experiment 4 in which sugar and nitrogen were normal. Remarkably the values of the NMRSE in experiment 4 were practically the same for both models.

Figure 3 shows the trajectories for all four candidate models and the corresponding experimental data for external compounds: YAN and glucose consumption plus ethanol production, as measured in three out of five experiments. The figure concentrates in those three experiments for which the differences between the best and the worst models are higher (namely Exp1, Exp2 and Exp5).

Results show that all models can explain the data with *R*^2^ > 0.9 for most cases. Different models presented different performance; in general, *M*_*P*_ and *M*_*X*_ could successfully recover all data with *R*^2^ > 0.96. *M*_*M*_ and *M*_*C*_ showed higher discrepancies with data. Remarkably *M*_*C*_ was the best to recover the final glucose consumption and ethanol production in experiment 5, which corresponds to a stuck fermentation.

## DISCUSSION

In this study, we proposed tw°Candidate models to explain nitrogen-limited fermentations of *S. cerevisiae*. We proposed two modifications to the standard modelling approaches: 1) the biomass growth accounts for protein and carbohydrates, and 2) fermentation rate is proportional to the total protein content. We reconciled candidate models to previously published data.

Our results revealed that Monod law is insufficient to explain biomass formation kinetics in nitrogen-limited fermentations of *S. cerevisisae*. This is so because, during exponential growth, the biomass composition seems to remain unaltered with mRNA and protein comprising an important percentage of the biomass. However, this is not the case during secondary growth phase where *µ*_1_ fades and *µ*_2_ increases, as shown in (13) data.

Several recent studies on the modelling of alcoholic fermentation of *S. cerevisisae* relied on logistic growth (20, 12), Monod or the use of kinetic constraints coupled to a stoichiometric model (which in practice holds similar results to Monod).

Logistic growth is well suited to explain cell numbers in nitrogen-limited fermentations. However, in light of the current results, we argue the same might not be valid in the case of biomass formation. A logistic curve model might not adequately describe the delay during the transition from exponential growth (*µ*_*N*_ >> *µ*_*C*_) t°Carbohydrate accumulation (*µ*_*C*_ >> *µ*_*N*_) and the different dynamics between *µ*_*N*_ and *µ*_*C*_ observed in cell mass data.

From a practical point of view, nitrogen-limited Monod models seem to explain biomass formation until the nitrogen source is depleted but struggle to explain later stages. The models proposed by Cramer et al. (8), Coleman et al. (9) distinguished active from inactive cell mass and used the methylene blue method to detect metabolic activity. Their experimental data shows that total and active biomass are practically equivalent until depletion of ammonia. After that, total biomass increases while active biomass stays stable or decreases. Because the model included Michaelis-Menten type expressions without ethanol inhibition, the excessive burden is put in the biomass decay - considered to be ethanol dependent. The authors observed that their model showed some discrepancies in the fits for cell mass. These discrepancies were more relevant at lower temperatures, for which the model predicted that cell activity decreased more rapidly than shown by the data. The models proposed in this work, outperformed Coleman et al. (9) results, but for the prediction of glucose and ethanol consumption in stuck fermentations. Our model could be further refined by introducing the concept of cellular decay to improve the accuracy in the prediction of stuck fermentations.

Malherbe et al. (20) developed a fermentation model for enological conditions that accounted for different temperatures and initial nitrogen concentrations. These authors estimated the number of hexose transporters present in the yeast cells as a function of time, temperature and nitrogen. The model represents glucose consumption data extremely well. However, it should be noted that it comes at the cost of a heavily parameterised and nonlinear hexose transport function that also includes inhibition by substrate and ethanol. Nevertheless, the notion that hexose transporters vary with nitrogen content along the fermentation is likely to find some overlap with our simpler model for glucose transport.

Pizarro et al. (17) included the effect of ethanol inhibition on sugar transport and the existence of different hexose transporters with different maximum transport rates and substrate affinities. From a qualitative point of view, this model corresponded to a significant improvement over past models. Noticeably, the hexose transport functions considered the role of the initial nitrogen concentration and temperature. Additionally, Varela et al. (16) showed that the protein content of biomass is heavily dependent on the initial nitrogen concentration which supports the notion that hexose transport is likely to be better described by total protein content rather than biomass *per se*. Our model would support this hypothesis. Results obtained by *M*_*P*_ model are superior to those obtained with the *M*_*X*_ model, at least, for those cases in which initial nitrogen is restricted.

Here we proposed an alternative modelling strategy which using a minimal number of parameters, that can be estimated from easily measured data, provides a considerable improvement in the description of nitrogen-limited fermentations as compared to previously published models. Other mechanisms, such as the role of ethanol in cellular decay, can be easily incorporated. The model brings a new possibility to anticipate, and thus control, problematic fermentations.

## MATERIALS AND METHODS

### Data

We considered data-sets from five different experiments as published in the literature. The first experiment (Exp1) data set was obtained from Schulze et al. (13). The authors conducted a nitrogen-limited (165 mg NH_3_ and 50g of glucose) batch fermentation while measuring biomass, protein content of biomass along with several extracellular metabolites (glycerol, ethanol and ammonia). We als°Considered four datasets from Coleman et al. (9). The authors performed various experiments at different conditions. In this work we denote Exp2 as that performed at 11*°C*, with high-sugar and low-nitrogen; Exp3, the one performed at 15*°C*, with normal sugar and low-nitrogen; Exp4, the one performed at 30*°C* with normal sugar and normal nitrogen; and Exp5, the one performed at 35*°C* with high-sugar and low-nitrogen.

### Model building

The proposed model is composed of six ordinary differential equations (as presented in the section Results). Solutions for the system are of type:

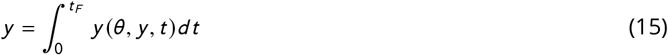

where *y* is the solution of the ODE system and corresponds to the dynamics of the relevant state variables - biomass, carbohydrates, protein, glucose, YAN, ethanol, glycerol-between the beginning and the end of the experiment (*t*_*f*_); *y* represents the states time derivative.

### Structural identifiability analysis

The proposed model depends on fourteen unknown parameters ***θ*** - namely 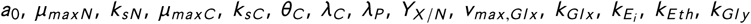 - to be estimated from the experimental data. The proposed model introduces the description of the total protein content. However for some experimental set-ups protein was not measured. Therefore we performed a structural identifiability analysis ((29, 30)) to assess if all parameters of the model can be estimated even if protein measurements are not available.

We performed the analysis using GenSSI2 toolbox (31). The toolbox uses symbolic manipulation to implement the so-called generating series of the model and t°Compute identifiability *tableaus*.

### Parameter estimation

The aim of parameter estimation is t°Compute the unknown parameters - growth related constants and kinetic parameters - that minimize the weighted least squares function which provides a measure of the distance between data and model predictions (**?**). In the absence of error estimates for the experimental data a fixed value for each state, the maximum experimental data, was assumed in order to normalize the least squares residuals:

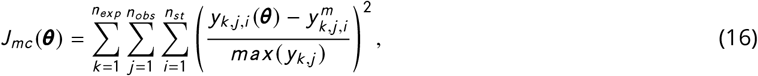

where *n*_*exp*_, *n*_*obs*_ and *n*_*st*_ are, respectively, the number of experiments, observables, and sampling times. 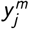 represents each of the measured quantities and *y*_*j*_ (***θ***) corresponds to model predicted values.

In our particular case, we used five different experiments: the first, taken from (13), and second to fifth, taken from (9). All experiments provided time series data of the biomass (*X*), glucose uptake (*Gl x*), amonia consumption (*N*) and ethanol production *E th*.

Parameters were estimated by solving a nonlinear optimization problem to find the unknown parameter values (***θ***) to minimize *J*_*mc*_ (***θ***), subject to the system dynamics - the model- and parameter bounds (32).

To avoid the risk of premature convergence by the optimization routine while searching for the optimal parameter set, the parameter estimation procedure was repeated 10 times for each candidate model starting from different initial guesses.

### Model selection

Models were compared in terms of the Akaike’s information criterion (AIC). AIC compares multiple competing models (or working hypotheses) in those cases where no single model stands out as being the best. AIC is calculated using the number of data (*n*_*d*_), the number fitted parameters (*n*_*θ*_) and the sum of squares of the weighted residuals (WRSS). For the cases in which the number of data is low relative to the number of parameters (roughly *n*_*d*_ /*n*_*θ*_ < 40 for the most complex model), the corrected Akaike’s information criterion is computed as follows:

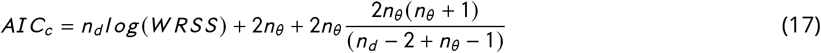

The models were ranked by *AIC*_*c*_, with the best approximating model being the one with the lowest *AIC*_*c*_ value. The minimum AIC value (AIC_*mi n*_) was used to re-scale the Akaike’s information criterion. The re-scaled value Δ = |*AIC*_*c*_ − *AIC*_*mi n*_ | was used to assess the relative merit of each model. Models such as Δ ≤ 2 have substantial support, models for which 4 ≤ Δ ≤ 7 have considerably less support and models with Δ > 10 have no support (33).

### Numerical methods and tools

The parameter estimation and model selection were implemented in the AMIGO2 toolbox (34). The system of ODEs was compiled to speed up calculations and solved using the CVODES solver (35), a variable-step, variable-order Adams-Bashforth-Moulton method. The parameter estimation problem was solved using a hybrid meta-heuristic, the enhanced scatter search method (eSS, (36)).

## SUPPLEMENTAL MATERIAL

All scripts and data required to reproduce results and figures can be accessed in https://sites.google.com/site/amigo2toolbox/examples.

## ACKNOWLEDGMENTS

This project has received funding from MCIU/AEI/FEDER, UE (grant reference RTI2018-093744-B-C33) and GAIN Xunta de Galicia (Contrato Programa and grant reference IN607B 2020-03).

